# A Combined Evidence Approach to Prioritize Nipah Virus Inhibitors

**DOI:** 10.1101/2020.03.12.977918

**Authors:** Nishi Kumari, Ayush Upadhyay, Kishan Kalia, Rakesh Kumar, Kanika Tuteja, Priyanka Rani Paul, Eugenia Covernton, Tina Sharma, Vinod Scaria, Anshu Bhardwaj

## Abstract

Nipah Virus (NiV) came into limelight due to an outbreak in Kerala, India. NiV causes severe disease and death in people with over 75% case fatality rate. It is a public health concern and has the potential to become a global pandemic. Lack of treatment has forced the containment methods to be restricted to isolation and surveillance. WHO’s ‘R&D Blueprint list of priority diseases’ (2018) indicates that there is an urgent need for accelerated research & development for addressing NiV. In the quest for druglike NiV inhibitors (NVIs) a thorough literature search followed by systematic data curation was conducted. Rigorous data analysis was done with curated NVIs for prioritizing druglike compounds. For the same, more than 1800 descriptors of NVIs were computed and comparative analysis was performed with the FDA approved small molecules and antivirals. These compounds were further evaluated through PAINS filter to study their toxicity profile. Simultaneously, compounds were also prioritized based on the robustness of the assays through which they were identified. Our efforts lead to the creation of a well-curated structured knowledgebase of 182 NVIs with 98 small molecule inhibitors. The reported IC50/EC50 values for some of these inhibitors are in the nanomolar range – as low as 0.47 nM. In order to prioritize these inhibitors, we performed several tests and applied filters to identify drug-like non-toxic compounds. Of 98, a few compounds passed DruLito & PAINS filters exhibiting drug-like properties and were also prioritized in an independent screen based only the assay robustness. The NVIs have diverse structural features and offer a wide spectrum of ways in which they can be developed further as druglike molecules. We report a knowledgebase for furthering the development of NVIs. The platform has a diverse set of 98 NVIs of which a few have been prioritized based on a combined evidence strategy. The platform has the provision to submit new inhibitors as and when reported by the community for further enhancement of NiV inhibitor landscape.

## Introduction

Nipah is an infectious negative-sense single-stranded RNA virus which belongs to the genus henipavirus and family Paramyxoviridae [1]. It is a pleomorphic enveloped virus with a particle size ranging from 40 to 1,900nm [2]. While fruit bats are thought to be the natural reservoirs of the virus, they are also able to spread to humans and some other species [3]. There are two major genetic lineages of the virus which are known to infect humans i.e, NIV Malaysia (NIV_M_) and NIV Bangladesh (NIV_B_) [2].

The first outbreak of Nipah virus infection erupted between September 29,1998, and April 4, 1999, when cases of febrile encephalitis in the suburb of Ipoh, Perak, Southern Peninsular Malaysia were reported to the Malaysian Ministry of Health (MOH) [4]. It was traced back to cross-species transmission of the virus from pigs to humans [4,5]. In 2001 another strain of the virus was reported to cause an outbreak in Meherpur, Bangladesh. Since then outbreaks have been recorded almost annually in the country and surrounding regions such as Siliguri, India [6,7]. Most of the outbreaks have been observed to have a mortality rate ranging from 50-100% [8]. Although direct contact with the infected bat’s contaminants could cause transmission of the virus, direct human-to-human transmission has also been observed in the case of healthcare professionals and relatives of infected patients [9,10]. More recently, the virus has garnered attention due to the first-ever outbreak in southern India during May of 2018 where 19 cases of NiV were reported in Kozhikode, Kerala. In the subsequent year, 2019, another case was reported in Ernakulam district of the state [11,12].

Nipah virus causes cell-to-cell fusion in the host thus forming multinucleated cells called syncytia. This allows the virus to spread despite the absence of viral budding and greatly influences its pathogenicity [13,14]. The incubation period of the virus ranges from 5 to 14 days (https://www.cdc.gov/vhf/nipah/pdf/factsheet.pdf). Primarily the virus infects the central nervous system but the NIV_B_ strain has been shown to have significant respiratory involvement [2]. High mutation rate, human to human transmission, survival time of up to 4 days under various environmental conditions and lack of treatment makes it a potential bioterrorism agent [15,16].

Currently, there is no known treatment for the disease. Broad-spectrum antiviral ribavirin has shown contradictory results with even the most optimistic ones significantly below 50% in causing reduction of the mortality rate [17,18,19]. Favipiravir (T-705), a purine analogue antiviral, is reported to provide protection to upto 14 days in the Syrian hamster model challenged with a lethal dose of Nipah virus [20]. More recently, experimental drug remdesivir has demonstrated complete protection to four African green monkeys who received a lethal dose of Nipah virus. Studies are planned to assess the efficacy of the treatment over time [21]. Despite being a lethal pathogen there is not a single drug under clinical trial against this virus. Due to several challenges in discovery and development of NVIs, along with lack of therapeutic intervention and the pathogenic propensity of the virus, Nipah infections are enlisted amongst WHO’s list of priority diseases [22].

As mentioned above, various groups have made efforts over the years to identify Nipah virus inhibitors using a variety of viral assays. Most of them have used syncytia formation or titre reduction assays to identify Nipah virus inhibitors. Some of the earlier studies report nipah inhibitors based on screening of large libraries within BSL-4 settings to identify compounds that inhibited the virus-induced cytopathic effect. Three sulfonamide compounds with low EC50 values are reported [23] from these screens. More recently, Bimolecular Multicellular Complementation assays have been developed which can be used to identify potential anti-nipah inhibitors by qualitatively and quantitatively investigating the formation of a syncytium [24]. Also reported recently is a humanized monoclonal antibody for cross-neutralizing NiV and HeV. Cryo-electron microscopy followed by fusion studies demonstrated that the antibody binds to a prefusion-specific quaternary epitope which is conserved in NiV F and HeV F glycoproteins. The binding prevents membrane fusion and viral entry and clearly indicates the importance of the HNV prefusion F conformation for eliciting a robust immune response [25].

Despite reports of nipah virus inhibitors, no systematic evaluation of the reported compounds is performed. A few antiviral inhibitor databases and studies exist, namely, The Influenza Research Database (IRD) [26][27], The Virus Pathogen Database and Analysis Resource (ViPR) [28], enamine library (https://enamine.net/hit-finding/focused-libraries/view-all/antiviral-library), etc. They contain very limited or no information with the focus on Nipah virus. The ones with focus on Nipah virus inhibitors are either scattered datasets or curated versions with limited or no evaluation of the anti-Nipah compounds as starting points for drug discovery and development of anti-Nipah inhibitor(s) [29].

Towards this, we developed a combined evidence based strategy to systematically evaluate Nipah virus inhibitors. With this objective, we curated the anti-Nipah compounds using a crowdsourcing model and the data were made publicly available at – http://vinodscaria.rnabiology.org/nipah since the inception of the project (May 2018). Our two pronged strategy comprised of (i) prioritization of NVIs based on the robustness of the assays which were used to identify them and (ii) estimation of several drug likeness parameters for the NVIs. The drug likeness analysis includes diversity analysis, similarity to FDA approved compounds (as drug repurposing candidates) or reported antivirals (to identify new scaffolds), profiling based on PAINS and DruLito filters and comparison to ZINC lead like libraries. The data curated from literature along with all these analyses is made available on a web-based platform which also allows submission and assessment of new molecules reported as NVIs.

## Material and Methods

### 1.1 Data source and structure

PubMed (https://www.ncbi.nlm.nih.gov/pubmed) and Google (https://www.google.com/) were the two main sources considered for collecting the inhibitors that showed activity against Nipah virus. Research papers relevant to the study were downloaded using the query as ‘Nipah’ and those with the keyword in their abstract/title were shortlisted. At the time of curation this keyword listed 838 papers which were considered for data-mining. This step was followed by a scoring system wherein papers that had confirmed information regarding the inhibitors against Nipah virus in their abstracts were assigned a score of ‘10’, those that may or may not have information regarding the same were given a score of ‘5’, those with information on peptides used as inhibitors were assigned a score of ‘4’ and rest with no relevant information were given a score of ‘0’. Papers with score values of 10, 5 and 4 were considered for further study and all of them were manually curated to create a Nipah Virus Inhibitor Knowledgebase (NVIK) sheet which is used by all the curators to organize data based on a predefined data structure shown below in Table 1.

**Table 1:** Data structure for NVIK

|  |  |
| --- | --- |
| ID - Internal ID for Nipah Virus Inhibitor Knowledgebase | Target |
| Inhibitor Name - as reported in literature | Assay type |
| PubChemID / DrugBank ID / ChemSpider ID | Assay description |
| SMILES (acc. to the structure in PubChem) | Assay source |
| Molecular weight | Assay cell type |
| IC50 / EC50 in [uM & nM] | *in vivo* testing performed |
| Reference (PMID/DOI) | Annotator Email |

For the structure of the inhibitors, MarvinSketch 5.11.4 software from ChemAxon (https://chemaxon.com/) was used and the corresponding ChemSpider (http://www.chemspider.com/) and PubChem (https://pubchem.ncbi.nlm.nih.gov/) Ids were fetched. These structures were either taken directly from the original paper or the references mentioned in it. Else they were retrieved directly from the company which sourced the compounds initially in the original paper. Repurposed drugs were given DrugBank Id. All inhibitors were given a unique Id. EC50/ IC50 values are mentioned in nM & uM units.

### 1.2 Data quality check

In order to ensure data quality, a separate team of curators re-performed the curation independent of the initial team. Errors are marked in ‘Red’ in the curation sheet and was then marked ‘Green’ once the data is verified by both the teams. Data curation and quality check was done manually and the data is stored in NVIK-Open resource Sheet which served as the crowdsourcing platform. Each entry was tagged with its contributor’s email Id.

### 1.3 Platform architecture

The Nipah Virus Inhibitor Knowledgebase (NVIK) is developed using the open source LAMP (Linux-Apache-Mysql-PHP) server technology. We have also used PHP, HTML, JavaScript, AJAX and CSS. The web-interface has provision for text, structure and complex query based search and other options for browsing.

#### Utility

##### 2 User interface

NVIK is designed as a platform to browse, search and submit NVIs. The search tools are described below. A submission form has been provided that can be used to submit new compounds in the database. The database also lists database statistics along with the physicochemical properties of the compound.

### 2.1 Search tools

In the NVIK web interface, three search options are available, namely, ‘Simple Search’, ‘Query Builder’ and ‘Structure Search’. The Simple Search option is to retrieve information using a text-based query on all/selected fields of the database. User can selectively display the searched records. Overall, the database contains – unique fields as shown in Figure 1.

**Figure 1:**
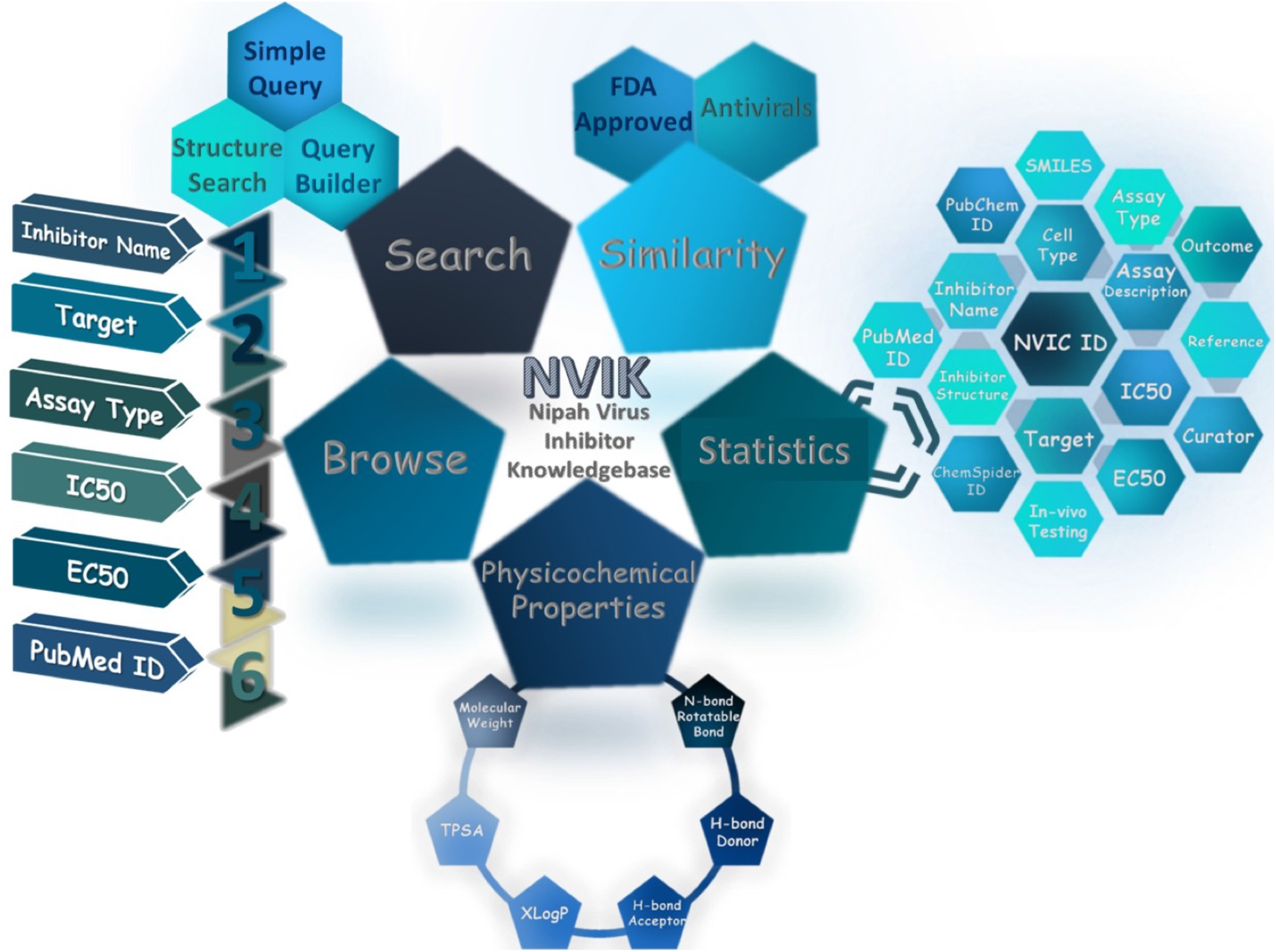
NVIK architecture illustrating various fields with ‘Search’ options, ‘Browse by’ options and structural ‘similarity’ options.

NVIK’s query builder option enables search using complex queries with the help of logical operators such as AND & OR. With these options, the user can retrieve targeted results such as compounds tested in specific cell lines with different IC50 values.

An extensive ‘Structure Search’ interface is also available in NVIK. The user can query using SMILES, SDF and MOL formats. In addition, the users can also draw a compound in JSME browser that may be searched in NVIK.

### 2.2 Browsing interface

Data on NVIs can be browsed for physicochemical properties like molecular weight, XLogP, polar surface area (PSA), hydrogen bond donor (HBD), hydrogen bond acceptor (HBA) and the number of rotatable bonds (http://bioinfo.imtech.res.in/anshu/nipah/physico_chem.php). The database can also be browsed from major fields as shown in Figure 1.

#### 3 Tools used for compound analysis and clustering

Near neighbours (NNs) in NVIK are identified using NNeib (version 5.12.0) (https://docs.chemaxon.com/display/docs/Jarvis-Patrick+clustering). The clusters of NVIs have been obtained using Jarp (version 6.0.2) (https://docs.chemaxon.com/display/docs/Jarvis-Patrick+clustering). which is based on the dissimilarity threshold (1-Tanimoto coefficient). A threshold of 0.15 is used as the dissimilarity coefficient. A total of 1875 descriptors were calculated using PaDel software [30]. To prioritize drug-like compounds, Drulito (http://www.niper.gov.in/pi_dev_tools/DruLiToWeb/DruLiTo_index.html) and PAINS filters were applied. Clustering is also performed with FDA approved compounds and the results are reported with known side-effects and predicted rat acute toxicity of NVIs similar to FDA approved compounds (http://bioinfo.imtech.res.in/anshu/nipah/fda_app.php).

##### Visualization tools

We have customized a CIRCOS [31] (http://circos.ca/) in one of our earlier studies on phytomolecules in mycobacteria [32]. We utilized a similar approach to represent the relationship among physicochemical properties of NVIs. In customized CIRCOS, 98 inhibitors are defined with a band having a width of 1000. This baseline is used to represent cyclic and aliphatic compounds. Appropriate modifications were then applied to generate further histogram files. To view the the compounds clusters, a compound network diagram (CND) was made using Cytoscape (http://www.cytoscape.org/).

## Results & Discussion

### NVIK statistics

At present, NVIK has 182 entries. The focus of the study was to evaluate small molecule inhibitors which were 125 entries as the rest 57 were either peptide or did not have complete information. Of these 125 entries, 98 are unique nipah inhibitors as for a few compounds more than one assay has been curated from the literature. Of these, EC50/IC50 values could be obtained for 70 compounds and structures are available for all 98 inhibitors. The EC50/IC50 values range from 0.25 nM to 7562710 nM for 182 entries and 0.47 nM to 7562710 nM for 125 small molecule entries. Some of these compounds are available in DrugBank and are predicted to have anti-nipah properties. The NVIK compounds are also mapped to Pubchem/Chemspider IDs wherever possible.

### Physicochemical properties and diversity in NVIKs

The overall distribution of important physicochemical properties of NVIs is depicted in Figure 2. It can be observed that most compounds in NVIK are cyclic in nature. Clustering of 98 NVIs resulted in 90 clusters of which 84 clusters (93.3% of NVIs) were singletons while remaining 14 clusters (15.5% of NVIs) were non-singletons. This illustrates that the NVIK library is chemically diverse. These clusters are represented as compound network diagrams made using Cytoscape. (http://bioinfo.imtech.res.in/anshu/nipah/circos.php) and the clusters are also reported in CIRCOS.

**Figure 2:**
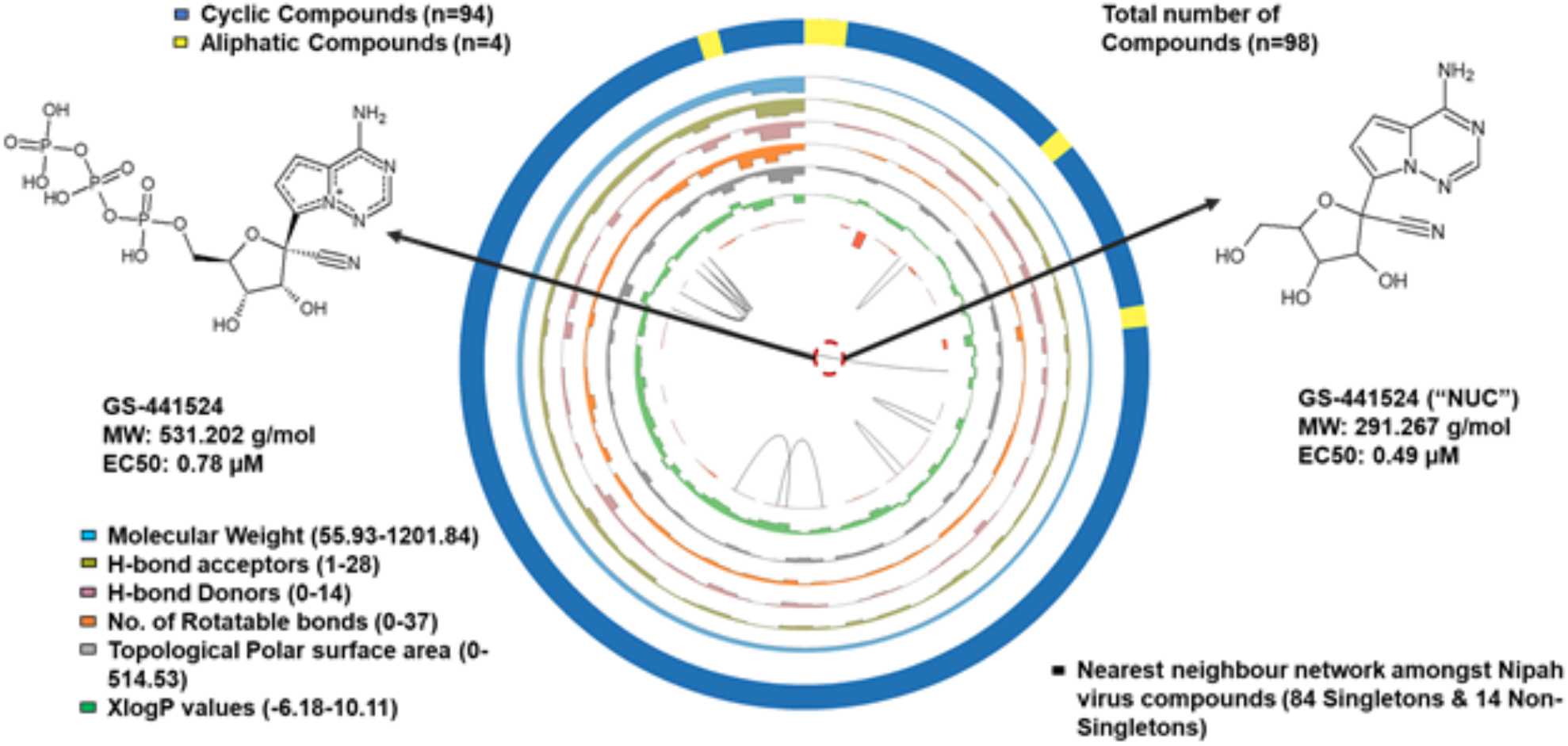
CIRCOS Plot illustrating the overall distribution of chemical and physicochemical properties of 98 NVIK small molecules. The diagram consists of 8 rings with the outermost ring categorizing the compounds as aliphatic or cyclic depending on their chemical nature. The subsequent rings illustrate the six graphs of calculated physicochemical properties. All compounds in the diagram are arranged in the increasing order of their molecular weight. The innermost ring represents the near neighbour network of NVIK compounds. Example of GS-441524 and GS-441524 (“NUC”) is shown. While the two compounds are structurally similar, the latter displays a better antiviral activity. It may be suspected that the slightest differences in the structure are responsible for this effect thus enabling derivation of pharmacophore features.

The innermost graph in Figure 2 represents the near neighbours among NVIs. An intriguing observation is that compounds with different molecular weight range are near neighbours. For example, GS-441524 (531.202 g/mol) is a near neighbour of GS-441524 (“NUC”) (291.267 g/mol). There may be common structural features in these compounds that are responsible for their activity against NiV (as highlighted). Therefore, this approach is useful in identifying potential pharmacophore features that are important in contributing to the antiviral properties of these compounds.

### Physicochemical property distribution of NVIs, FDA approved compounds and Antivirals

Comparison of NVIs is done with FDA approved compounds (2346) and antivirals obtained from Enamine library (4644). Enamine is a library of nucleoside like compounds (https://enamine.net/hit-finding/focused-libraries/view-all/antiviral-library). On the basis of physicochemical properties as suggested by Lipinski and Veber [33] it is observed that for the majority of NVIs, the distribution of all the six properties is similar to FDA approved drugs and also in some cases with Enamine compounds (Figure 3). Drugs with good oral availability, demonstrate TopoPSA ≤ 140 Å2 and number of rotatable bonds ≤ 10.

**Figure 3:**
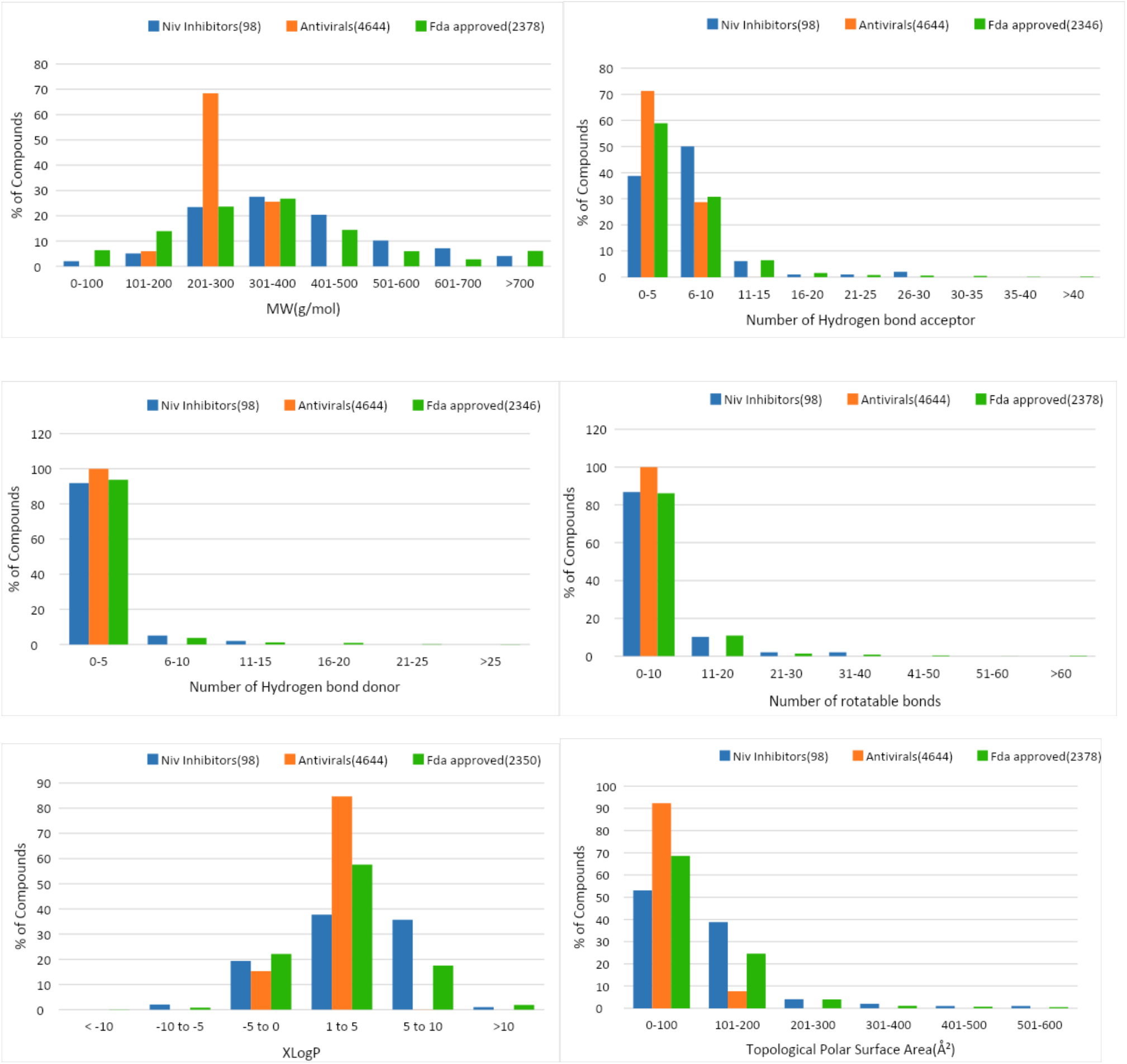
Physicochemical distribution of NVIs with respect to FDA approved and enamine compounds based on six properties. These properties were studied using the PaDel software. It can be observed that most NVIs have properties which overlap with FDA approved drugs or antiviral compounds.

### Structural similarity of NVIs with existing drugs

In total, there are 34 NVIs similar to 76 FDA approved drugsf. For e.g, 5’–deoxy-5’-methylthioadenosine (NVIC0097) has shown structural similarity against four FDA molecules, namely, Adenosine (a neurotransmitter), Vidarabine (active antiviral against herpes, vaccinia & varicella zoster virus), Fludarabine (a chemotherapy drug) and Ademetionine (used in the treatment of chronic liver diseases). In addition, side effects and predicted rat toxicity as reported in DrugBank is also listed for FDA approved compounds similar to NVIs (http://bioinfo.imtech.res.in/anshu/nipah/fda_app.php). This can aid in prioritization of NVIs with a potential safe profile as opposed to the ones which have reported side effects.

### Drug-likeness of NVIs

In order to prioritize druglike NVIs, DruLito is used which screens all compounds based on eight filters, namely, Lipinski’s rule, MDDR-like rule, Veber rule, Ghose filter, BBB rule, CMC-50 like rule, weighted and unweighted Quantitative Estimate of Drug-likeness [34][35][36]. As can be seen from Figure 4 only two of 91 compounds passed all filters other than CMC filter.

**Figure 4:**
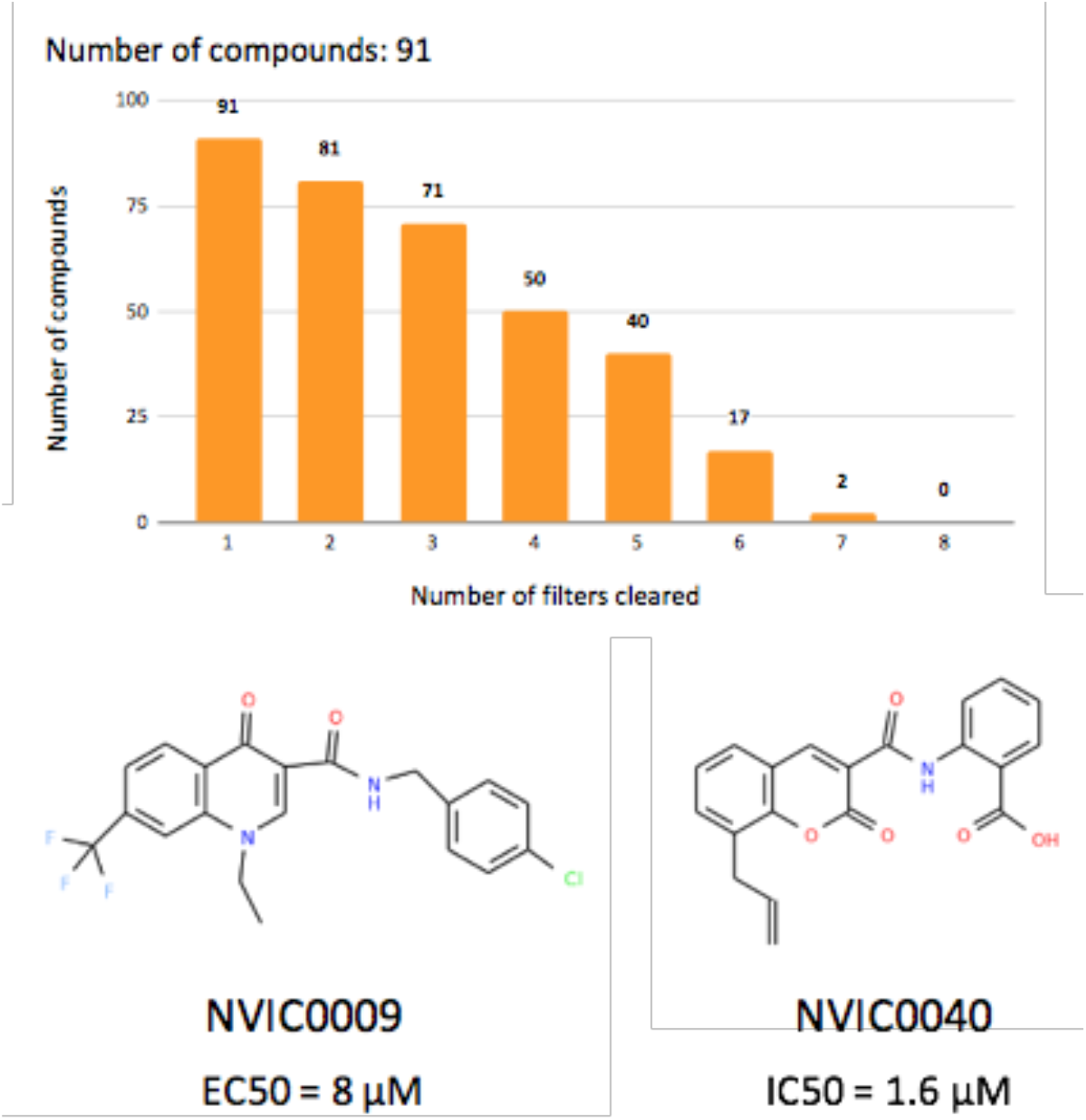
Drug-likeness study of NVIK compounds. (A) Compound distribution graph of 91 NVIs that can be filtered by DruLiTo (Drug-likeness Tool) filter with 2 compounds passing seven filters. More than half of the compounds pass at least four DruLito filters. (B) Structure of NVIK compounds crossing seven filters of DruLiTo: **NVIC0009**-N-(Cyclopropylmethyl)-1-ethyl-6-fluoro-4-oxo-1,4-N-(4-Chlorobenzyl)-1-ethyl-4-oxo-7-(trifluoromethyl)-1,4-dihydroquinoline-3-carboxamide and **NVIC0040** – 848-0115

### Combined evidence approach for prioritizing NVIs

In order to prioritize NVIs for further studies, we performed an independent evaluation of the compounds based on the robustness of the assays that were used to identify them. Based on this analysis, we prioritised 11 NVIs. Further analysis of these NVIs also indicated that they have physicochemical descriptor values which are closer to existing drug like FDA approved compounds. Our analysis of FDA approved compounds for several physicochemical parameters namely molecular weight, number of hydrogen bond donors, number of hydrogen bond acceptors, number of rotatable bonds, XlogP and topological polar surface area revealed that most prioritized NVIs have drug like features. As can be seen from Table 2, NVIC0025, NVIC0032 and NVIC0048 passed 6 DruLito filters and were accepted in FAF drugs analysis. Of the three compounds shortlisted NVIC0048 displayed a high degree of overlap with all six physicochemical parameter ranges. However NVIC0032 (9167 (2,2,2-trifluoro-N-[3-(N-naphthylcarbamoyl)(4,5,6,7-tetrahydrobenzo[β]thiophen-2-yl)]acetamide) exhibits better activity among the three and has considerable overlap with the physicochemical properties of FDA approved drugs. The scope of the current study was to perform a systematic evaluation of physicochemical properties of NVIs with respect to FDA approved drugs and antiviral libraries, understand the diversity of chemical scaffolds and correlate these computed properties with the experimentally reported bioactivity datasets for proposing a list of prioritized candidates. In order to simplify data access and further enhancement of data libraries, a web-based platform is designed for reporting new compounds, further analysis and prioritization of potential NVIs.

**Table 2 :** List of prioritized NVIs along with their biological activity, drug likeness profile and structural similarity to antivirals / FDA approved small molecular libraries. Most compounds were found to be dissimilar to FDA approved small molecule with the exception of bortezomib (NVIC0027) and chloroquine (NVIC0048). *minimum value has been reported **minimum effective concentration. None of these compounds show similarity to the Enamine antiviral library. #http://bioinfo.imtech.res.in/anshu/nipah/chembl_sim.php ##http://bioinfo.imtech.res.in/anshu/nipah/zinc_sim.php

| Compound ID | IC50[uM] | EC50[uM] | No. of DriLito filters followed | FaF Drugs4 |  |  | Similarity (Tanimoto Coefficient (TC)) |  |  |
| --- | --- | --- | --- | --- | --- | --- | --- | --- | --- |
|  |  |  |  | Accepted | Rejected | PAINS | FDA Approved (TC > 0.8) | No. of ChEMBL<sup>#</sup> ID's (TC > 0.85) | No. of ZINC Lead-like<sup>##</sup> ID's (TC > 0.85) |
| NVIC0002 | 0.029* | - | 1 |  | ✓ |  | X | 51 | 15 |
| NVIC0021 | - | 0.281±0.0448* | 3 | ✓ |  |  | X | X | 3 |
| NVIC0023 | 0.25** | - | 4 | ✓ |  |  | X | X | 6 |
| NVIC0024 | 0.149* | - | 5 |  | ✓ |  | X | 129 | 400 |
| NVIC0025 | 0.525 | - | 6 | ✓ |  |  | X | 3 | 6 |
| NVIC0026 | 0.218 | - | 5 |  |  | ✓ | X | 5 | 33 |
| NVIC0027 | 0.0027 | - | 5 |  |  | ✓ | DB00188 | 400 | 400 |
| NVIC0028 | 0.00047 | - | 1 | ✓ |  |  | X | 400 | 400 |
| NVIC0029 | 0.07 | - | 4 |  | ✓ |  | X | 400 | 400 |
| NVIC0032 | - | 0.0289±0.0102* | 6 | ✓ |  |  | X | 16 | 356 |
| NVIC0048 | 20.62 | - | 6 | ✓ |  |  | DB00608, DB01611 | X | 400 |

## Conclusions

NVIK is a unique effort to systematically map the chemical space for NVIs currently being pursued. The highlight of the approach is that prioritization of NVIs was performed not only by evaluating their druglike properties but also based on the robustness of the assays which were performed to identify them. Given that WHO’s ‘R&D Blueprint list of priority diseases’ (2018) indicates that there is an urgent need for accelerated research & development for addressing NiV, we believe that this platform may offer a centralized resource for the community to submit and access NVIs in a seamless manner. Our efforts have led to the creation of a well-curated structured knowledgebase of 182 NVIs with 98 small molecules. Based on a combined evidence strategy we have ranked the top 10 NVIs and have provided information on similar molecules from approved drugs, ChEMBL and ZINC libraries. We believe that the future of drug discovery of outbreak strains is in engaging wider scientific community in continous curation and analysis for these datasets. Not only for Nipah but these compounds may be utilized for their antiviral activity for other viruses too. Our effort has made available compound structure data that is readily available for these studies. The platform has the provision to submit new inhibitors as and when reported by the community for further enhancement of NiV inhibitor landscape.

## Availability and requirements

NVIK is freely accessible at: http://bioinfo.imtech.res.in/anshu/nipah/. To access all the features of NVIK to its optimum level, JavaScript and Java Runtime Environment (JRE) plugin must be enabled.

## Abbreviations

FDA: Food and Drug Administration
NVIs: Nipah Virus Inhibitors
Niv: Nipah Virus
WHO: World Health Organization
R&D: Research and Development
PAINS: Pan-assay interference compounds
IC50: Half maximal inhibitory concentration
EC50: half-maximal effective concentration
NVIK: Nipah Virus Inhibitor Knowledgebase
NVIC: Nipah Virus Inhibitor Collection
LAMP: Linux-Apache-Mysql-PHP
SMILES: SDF: structure data file
Nneib: Near Neighbour
TopoPSA: topological polar surface area
NADH: Nicotinamide adenine dinucleotide

## Author’s Contribution

NK, KT compiled data from the literature. NK, KT, PRP an AU curated the data and updated the field wherever data was incomplete. NK and PRP curated chemical structures. KT, KK and RK developed the database, web-based tools and the interface of NVIK with inputs from NK and AB. NK performed compound analysis. TS and NK plot visualization plots, CIRCOS and CND. EC performed assay robustness analysis and ranking, NK, KT, PRP, AU and AB drafted the manuscript. VS and AB conceived and designed the project and refined the manuscript. VS and AB defined the SOP for data collection and also the data structure. The manuscript has been read and approved by all authors.

## Acknowledgement

This study is an outcome of a community effort and we thank all the members who participated.

